# Benchmarking of novel green fluorescent proteins for the quantification of protein oligomerization in living cells

**DOI:** 10.1101/2023.02.07.527542

**Authors:** Annett Petrich, Amit Koikkarah Aji, Valentin Dunsing, Salvatore Chiantia

## Abstract

Protein-protein-interactions play an important role in several cellular functions. Quantitative non-invasive techniques are applied in living cells to evaluate such interactions, thereby providing a broader understanding of complex biological processes. Fluorescence fluctuation spectroscopy describes a group of quantitative microscopy approaches for the characterization of molecular interactions at single cell resolution. Through the obtained molecular brightness, it is possible to determine the oligomeric state of proteins. This is usually achieved by fusing fluorescent proteins (FPs) to the protein of interest. Recently, the number of novel green FPs has increased, with consequent improvements to the quality of fluctuation-based measurements. The photophysical behavior of FPs is influenced by multiple factors (including photobleaching, protonation-induced “blinking” and long-lived dark states). Assessing these factors is critical for selecting the appropriate fluorescent tag for live cell imaging applications. In this work, we focus on novel green FPs that are extensively used in live cell imaging. A systematic performance comparison of several green FPs in living cells under different pH conditions using Number & Brightness (N&B) analysis and scanning fluorescence correlation spectroscopy was performed. Our results show that the new FP Gamillus exhibits higher brightness at the cost of lower photostability and fluorescence probability (*pf*), especially at lower pH. mGreenLantern, on the other hand, thanks to a very high *pf*, is best suited for multimerization quantification at neutral pH. At lower pH, mEGFP remains apparently the best choice for multimerization investigation. These guidelines provide the information needed to plan quantitative fluorescence microscopy involving these FPs, both for general imaging or for Protein-protein-interactions quantification via fluorescence fluctuation-based methods.

## Introduction

A multitude of cellular processes, such as biomolecule transport, ion channel activity, cell-cell adhesion and communication are regulated by protein-protein-interactions (PPIs) (1–3). “Classical” bulk biochemical *in vitro* methods that are used to quantify PPIs (e.g., co-immunoprecipitation (co-IP), pull-down assays and western blotting) cannot be used to obtain information about intracellular protein distribution in live-cell samples or to monitor the effects of variations in concentrations between different cells (4, 5). Conventional optical microscopy can visualize the localization of proteins, but its resolution is limited (4, 6). More complex approaches, such as fluorescence fluctuation spectroscopy (FFS), can assess the interactions between molecules in complexes and obtain insights into cellular pathways and assembly processes (4–8). FFS provides information about dynamics through the analysis of signal fluctuations from fluorescently labeled molecules (6, 7, 9). Additionally, the magnitude of such fluctuations can be used to derive quantitative information about the multimerization state (i.e., number of monomers in a multimer) of the protein of interest (7–10).

A common strategy to investigate PPIs *in cellula* via FFS is the fusion of a fluorescent protein (FP) to the protein of interest (7–9, 11). Comparison between the brightness of protein multimers tagged with FPs and the brightness of a monomeric reference allows the quantification of FP monomers in the complex and, thus, the oligomerization state of the protein of interest (5, 7, 12). A major problem for several FFS applications that rely on FPs, though, is the presence of non-emitting “dark” proteins, which can be quantified through the so-called fluorescence probability (*pf*) (7, 13). It is sometime assumed that FPs have a *pf* value of 1 meaning that, for example, a trimer containing three FPs emits in average a three-fold higher signal than a monomer labelled with one FP (5, 7). Instead, *pf* values of e.g. green emitting FPs have been reported in the range between 0.5 and 0.8 (1, 2, 7, 13–19). Recently, we have systematically quantified the *pf* of several FPs, focusing mainly on proteins emitting in the red part of the visible spectrum (7). In summary, since high *pf* values are required for increased sensitivity, not all FPs are equally suitable for oligomerization studies (7). Because of the presence of non-emitting FPs (i.e., *pf* lower than 1), FFS approaches might underestimate the amount of FPs and therefore, the oligomeric state of the protein of interest.

Some of the most used FPs are the green fluorescent protein (GFP) and its mutants, such as the monomeric enhanced GFP (mEGFP) (5, 20). This FP has a high quantum yield (QY), enhanced photostability, and minimal interference with the cellular machinery (20). Of note, several variants have been engineered over the past few years (e.g. mNeonGreen (mNG), mGreenLantern (mGL), and Gamillus) in an effort to optimize molecular brightness, folding efficiency, photostability, and pH stability of the fluorescent probe (Table 1) (21–24).

**Table 1:** Photophysical characteristics of the fluorescence proteins (FPs). QY, Quantum yield; norm., normalized.

| FP | $\lambda_{\text{ex}}$ [nm] | $\lambda_{\text{em}}$ [nm] | QY | Brightness* | cellular<br>norm. <I> | pK<sub>a</sub> | maturation<br>[min] | Reference |
| --- | --- | --- | --- | --- | --- | --- | --- | --- |
| mEGFP | 488 | 507 | 0.71 | 38.0 | 1.0<sup>#,†</sup> | 6.0 | 28.0 | (21) |
| mNG | 506 | 516 | 0.78 | 90.0 | 3.3<sup>#</sup> | 5.7 | 18.0 | (21) |
| mGL | 503 | 514 | 0.72 | 74.0 | 6.3<sup>#</sup> | 5.6 | 14.0 | (21) |
| Gamillus | 504 | 519 | 0.90 | 74.7 | 0.5<sup>†</sup> | 3.4 | 8.0 | (24) |

\* (extinction coefficient × QY)/1,000, where QY and extinction coefficients are measured at the absorbance peak ( $\lambda_{\text{ex}} \pm 2$ nm)
<sup>#</sup> Average intensity in FP-expressing BE (2)-M17 human neuroblastoma cells, relative to EGFP, using the P2A quantitative co-expression system
<sup>†</sup> Average intensity in FP-expressing HeLa cells, relative to EGFP

In the context of complex experiments in living cells, a FP might encounter different environments and pH conditions (e.g. acidic pH in lysosomes, secretory granule and endosomes; neutral pH in the nucleus, cytoplasm and endoplasmic reticulum (ER); basic pH in mitochondria and a pH gradient in the Golgi network) (22, 25). Measurements of GFP at acidic pH have shown a decrease of the *pf* value, possibly due to an increase of proteins in the dark-state (22, 26–28). Consequently, it would be useful to systematically analyze the properties of the above-mentioned novel FPs, in order to perform an exact quantification of PPIs, also in different intra-cellular environments.

Here, we benchmarked the performance of novel green FPs (mGL and Gamillus) against the well-established mEGFP and mNG. The presence of non-fluorescent states and photostability under different pH conditions was measured using Number & Brightness (N&B) and scanning fluorescence correlation spectroscopy (sFCS) analysis (5). Our results indicate that some of the observed proteins are brighter, although unstable under longer/stronger illumination. Furthermore, we identify which proteins are more suitable for multimerization quantification, rather than “simple” imaging, also at different pH conditions.

## Materials and Methods

### Fluorescent protein constructs

A detailed description of the cloning procedure of all constructs is available in the Supporting Information. All plasmids generated for this work will be made available on Addgene (https://www.addgene.org/). A schematic overview of the intracellular localization of the constructs is provided Fig S1A and an overview of the linker sequences within the FP structures is provided in Table S1.

### Cell culture

Human embryonic kidney cells from the 293T line (HEK293T, CRL-3216^™^) and Chinese hamster ovary cells (CHO-K1, CCL-61^™^) were purchased from ATCC (Kielpin Lomianki, Poland). Both cell lines were cultured in phenol red-free Dulbecco’s modified Eagle’s medium (DMEM) with 10% fetal bovine serum, 2 mM L-glutamine, 100 U/mL penicillin, and 100 mg/mL streptomycin at 37 °C and 5% CO_2_. Cells were passaged every 2–3 days when they reached nearly 80% confluence in tissue culture flask, for 15 passages. All solutions, buffers, and media used for cell culture were purchased from PAN-Biotech (Aidenbach, Germany).

### Preparation for Microscopy Experiments

For microscopy experiments, 6 × 10^5^ (HEK293T) or 4 × 10^5^ (CHO-K1) cells were seeded in 35 mm dishes (CellVis, Mountain View, CA, USA) with optical glass bottom (#1.5 glass, 0.16–0.19 mm), 24 h before transfection. HEK293T cells are preferred for sFCS measurements since they are relatively thick and therefore suitable for sFCS based data acquisition perpendicular to the PM. CHO-K1 cells are rather flat and characterized by a large cytoplasmic volume and therefore suitable for N&B measurements in the cytoplasm. Cells were transfected 16–24 h prior to the experiment using 200 ng plasmid DNA per dish with Turbofect (Thermo Fisher Scientific, Waltham, MA, USA) according to the manufacturer’s instructions.

For measurements under different pH conditions, the culture medium was exchanged with buffer containing 140 mM NaCl, 2.5 mM KCl, 1.8 mM CaCl2, 1.0 mM MgCl2, and 20 mM HEPES with a pH value of 5.6, 7.4 or 9.2.

### Confocal microscopy system and setup calibration for fluorescence fluctuation spectroscopy

All measurements were performed on a Zeiss LSM780 system (Carl Zeiss Microscopy GmbH, Oberkochen, Germany) using a Plan-Apochromat 40×/1.2 Korr DIC M27 water immersion objective and a 32-channel GaAsP detector array. Samples were excited with a 488 nm Argon laser and the fluorescence signal was collected in a range of 498 to 606 nm in photon-counting mode after passing through a 488 nm dichroic mirror. For the spectral analysis under different pH conditions, fluorescence was detected in spectral channels of 8.9 nm width (23 channels between 491 nm and 690 nm). To decrease out-of-focus light, a pinhole with size corresponding to one airy unit was used. All measurements were performed at 22 ± 1°C.

The confocal volume was calibrated daily by performing a series of point FCS measurements with Alexa Fluor_®_ 488 (AF488, Thermo Fischer, Waltham, MA, USA) dissolved in water at 30 nM, at the same laser power and beam path used for N&B and sFCS measurements. Prior to that, the signal was optimized by adjusting the collar ring of the objective and the pinhole position to the maximal count rate for AF488. Then, five measurements at different locations were acquired, each consisting of 15 repetitions of 10 s, and the data was fitted using a three-dimensional diffusion model including a triplet contribution. The structure parameter *S* (defined as the ratio between the vertical and lateral dimension of the theoretical confocal ellipsoid) was typically around 5 to 9, and the diffusion time *τ_d_* around 30 to 35 μs.

### N&B measurements

N&B analysis was performed as previously described (7, 10) with few modifications: 128x 128 pixel images were acquired with pixel dimensions of 400 nm and a pixel dwell time of 50 μs. Image time-stacks of 105 scans were collected using the Zeiss Black ZEN software. The intensity time-stacks data were analyzed using a custom Matlab code (The MathWorks, Natick, MA, USA). The Matlab algorithm utilizes the equations from Digman *et al*. (11) for obtaining the molecular brightness and number as a function of pixel position. Bleaching and minor cell movements are partially corrected using a boxcar-filter with an 8-frame window applied pixel-wise, as previously described (7, 29, 30). Final brightness values were calculated by extrapolating the partial brightness values (i.e., calculated within each 8-frame window) to the earliest time point. Detector saturation leading to artefactual reduction in brightness was avoided by excluding pixels with photon-counting rates exceeding 1 MHz. A schematic overview of the N&B analysis is provided in Fig S1C.

### sFCS measurements

Scanning FCS experiments were performed as previously described (7, 31), with a modified acquisition mode. Briefly, a line scan of 256 × 1 pixels (pixel size ≈80 nm) were performed perpendicular to the membrane with 472.73 μs scan time. Typically, 400,000 lines were acquired (total scan time ≈3 min) in photon counting mode. Laser power values were typically between ≈1.5 μW (brightness analysis) and ≈6 μW (photobleaching analysis). Scanning data were exported as TIFF files, imported and analyzed in Matlab (The MathWorks, Natick, MA, USA) using a custom code as previously described (7, 31, 32). A schematic overview of the sFCS analysis is provided in Fig S1D. Matlab custom-written code is available from the corresponding author upon reasonable request.

### Brightness calibration and fluorophore maturation

The molecular brightness, i.e. the photon count rate per molecule, is used as a measure for the oligomeric state of protein complexes. This quantity is often based on the assumption that all fluorophores within an oligomer are fluorescent. However, FPs can undergo dark state transitions or be in a non-mature, non-fluorescent state (8). To quantify the amount of non-fluorescent FPs, we consider all these processes together in a single parameter, the apparent fluorescence probability (*pf*), i.e. the probability of a FP to emit a fluorescence signal. The determination of the *pf* value from apparent brightness values was performed as previously described (7, 32). Shortly, for each sample and each experiment day, the average brightness for a monomeric construct was determined from multiple cells (see e.g. Figs 2A and 3A). Then, measurements were performed in several cells expressing the dimer constructs (Fig S2) and, from each of these measurements and the average monomer brightness obtained before, one *pf* value was calculated using the formula *p_f_* = ^*Brightness_dimer_*^/_*Brightness_monomer_*_ −1 (7, 32). The final *pf* value was calculated as mean of such a set of measurements (see Figs 2B and 3B).

**Fig 1:**
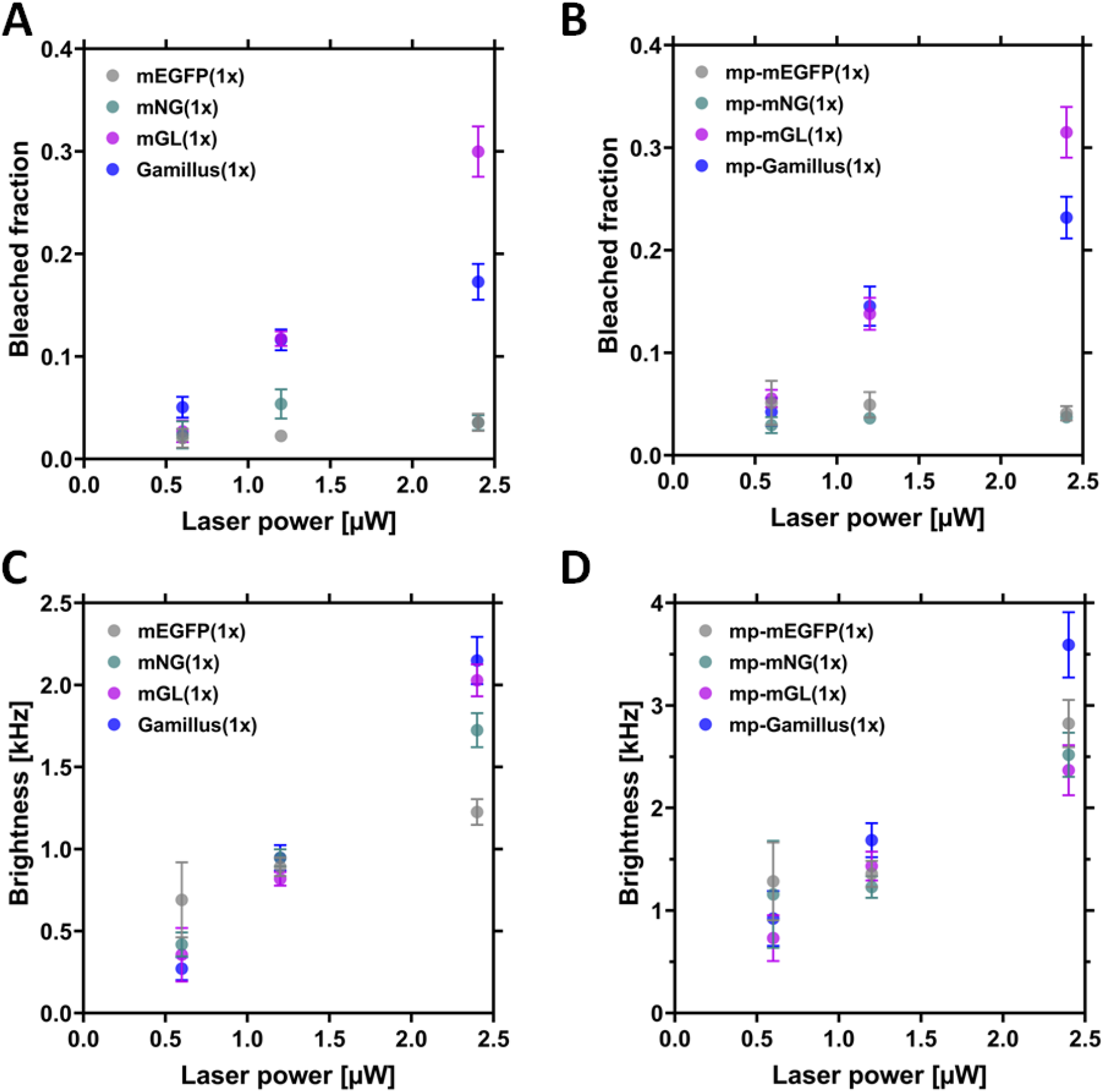
Comparison of the molecular brightness and bleached fraction for the monomeric FPs, as obtained from N&B measurements in CHO-K1 cells. CHO-K1 cells were transfected with plasmids coding for the monomeric cytosolic FPs (mEGFP(1x), mNeonGreen(1x) (here, called mNG(1x)), mGreenLantern(1x) (here called mGL(1x)) and Gamillus(1x)) or with membrane-anchored FPs. The latter are anchored to the inner leaflet of the PM via a myristoylated and palmitoylated (mp) peptide (mp-mEGFP(1x), mp-mNG(1x), mp-mGL(1x) or mp-Gamillus(1x)). N&B measurements were performed ≈16 h after transfection with a laser power of 0.6 μW, 1.2 μW and 2.4 μW. (A-B): Mean bleached fractions measured for cytosolic FPs (A) and for PM-anchored FPs (B), as a function of laser power. The bleached fraction indicates the amount of fluorescence signal lost after a N&B measurement. Error bars indicate the standard error of the mean (SEM). (C-D): Mean brightness values measured for cytosolic FPs (C) and for PM-anchored FPs (D), as a function of laser power. Error bars represent the SEM. Exact values and sample sizes are summarized in Table S2.

**Fig 2:**
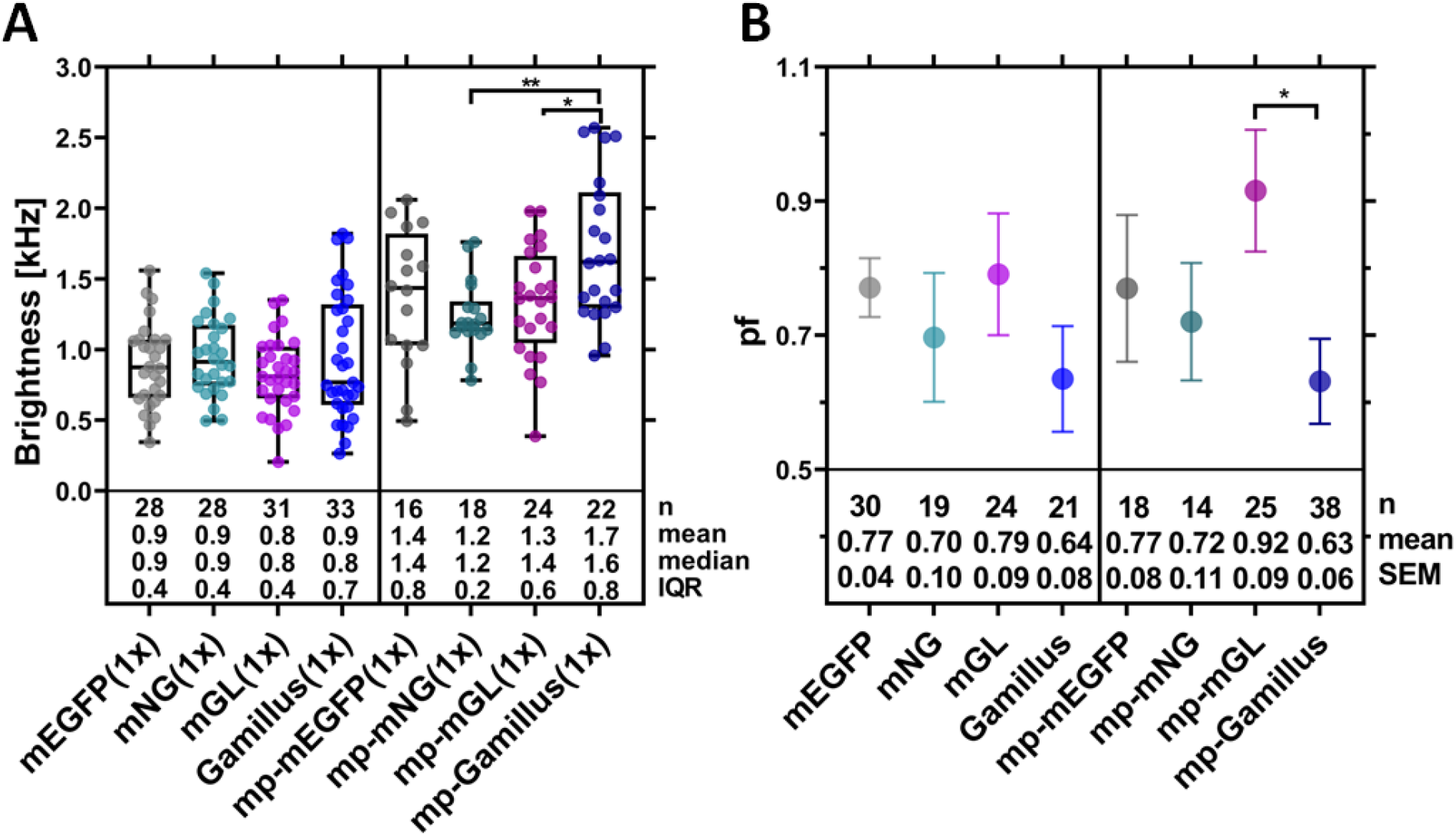
Comparison of brightness and fluorescence probability (*pf*) for the examined green FPs, as obtained from N&B measurements in CHO-K1 cells. N&B measurements were performed ≈16 h after transfection, using a laser power of 1.2 μW. (A): Box plot of the molecular brightness for the examined cytosolic and membrane-anchored FPs (i.e., mp-FP). Each point represents the average value measured in a single cells, from three independent experiments. Median values and whiskers ranging from minimum to maximum values are displayed. (B): Mean *pf* values calculated for the cytosolic and membrane-anchored FPs, using the brightness values measured for the corresponding FP dimers (Fig S2A). Data were collected from three independent experiments. The error bars represent the SEM. Sample size, mean/median, and interquartile range (IQR)/SEM are indicated below the graph. Statistical significance within both plots for selected sample pairs were determined using one-way ANOVA Tukey’s multiple comparison test (* p < 0.05, ** p < 0.005).

**Fig 3:**
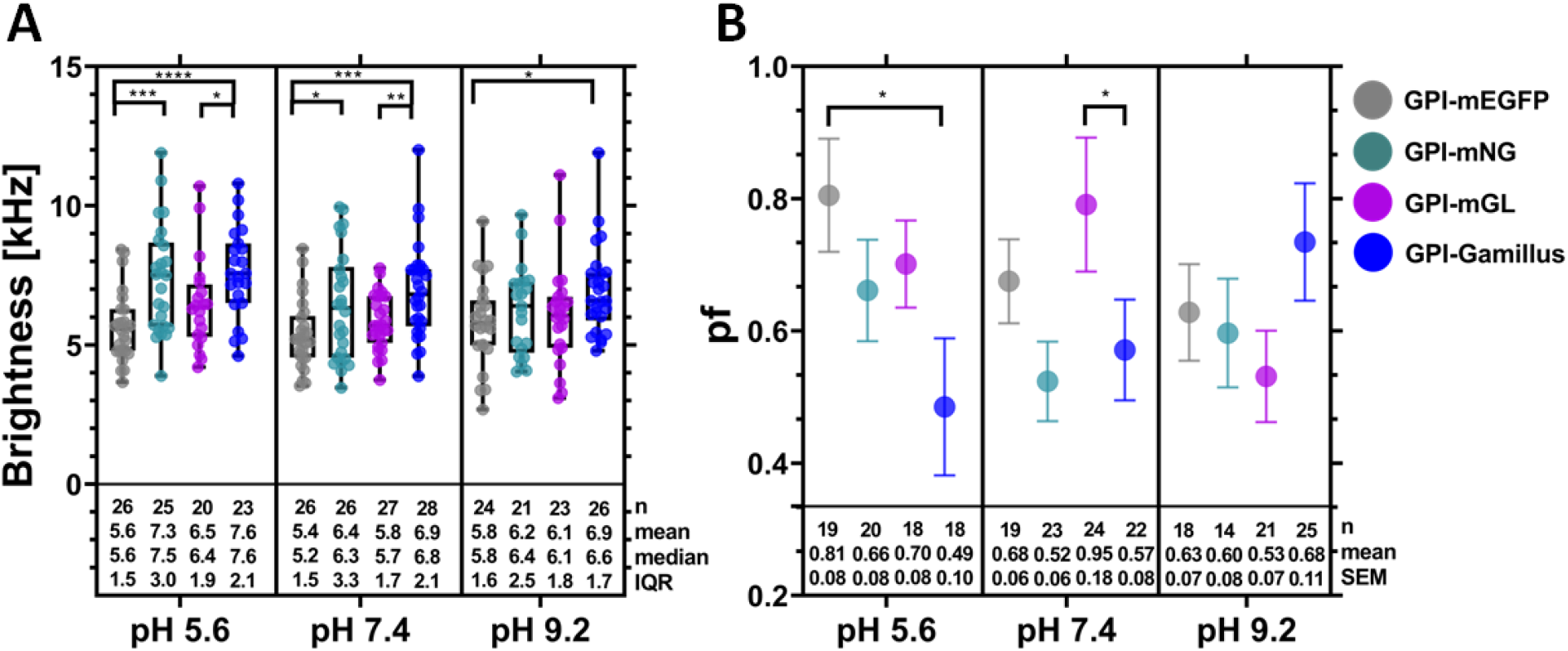
Comparison of brightness and *pf* values for the examined FPs, as obtained from scanning fluorescence correlation spectroscopy (sFCS) measurements in HEK293T cells, under different pH conditions (pH 5.6, pH 7.4, and pH 9.2). sFCS measurements were performed ≈16 h after transfection, using a laser power of 1.5 μW. (A): Box plots of the molecular brightness for the membrane-anchored FPs GPI-mEGFP, GPI-mNG, GPI-mGL and GPI-Gamillus, for three different pH conditions. Each point represents the average value measured in single cells, from three independent experiments. Median values and whiskers ranging from minimum to maximum values are displayed. (B): Average *pf* values calculated using the brightness values measured for the corresponding FP dimers (Fig S2B), for three different pH conditions. Data were collected from three independent experiments. The error bars represent the SEM. Statistical significance was determined for selected sample pairs in both plots using one-way ANOVA Fishers Least Significant Difference (LSD) multiple comparison test (* p < 0.05, ** p < 0.005, *** p < 0.0005, **** p < 0.0001).

### Statistical analysis

Data from at least three independent experiments were pooled and visualized using GraphPad Prism ver. 9.0.0 (GraphPad Software, LCC, San Diego, CA, USA). All resulting data were displayed as box plots with single data points corresponding to measurements in single cells or as mean ± standard error of the mean (SEM) plots. Median values and whiskers ranging from minimum to maximum values are displayed in the box plots. The mean, median, interquartile range (IQR) are indicated in each graph together with the sample size. Significance values are given in each graph and figure captions, respectively. Statistical significance was tested by using D’Agostino-Pearson normality test followed by one-way ANOVA analysis and the Tukey’s or Fishers Least Significant Difference (LSD) multiple comparisons test.

## Results

### mGL and Gamillus are less photostable than mEGFP or mNG

All monomeric FPs were evaluated for photostability at different laser powers using confocal microscopy, in the context of a typical N&B experiment performed in CHO-K1 cells. Such an approach simply consists of a time-series acquisition with continuous scanning over ca. two minutes. mGL and Gamillus displayed higher photobleaching, both in the cytosol and at the PM, in comparison to mEGFP and mNG (Fig 1A and 1B, Table S2). For example, ≈30% of the total mGL was bleached after a ca. two-minute of continuous scanning, using 2.4 μW excitation power. At the same time, N&B analysis provided the brightness of each FP, as a function of the laser power. Although an accurate analysis should be performed at an excitation power causing as low as possible bleaching (see next paragraph), Figs 1C and D confirm that the observed brightness predictably increases roughly linearly with the laser power. As expected, due to the detection geometry, a higher signal is observed for molecules restricted to the PM (33). Finally, Gamillus shows generally a higher brightness, as especially noticeable for measurements at the PM or at higher excitation powers.

### Gamillus exhibits high brightness but low fluorescence probability

To compare the brightness effectively observed in a typical confocal microscopy setup and, specifically, in the context of FFS experiments, we have performed in-depth N&B analysis of FP monomer and dimer constructs. In order to maintain bleaching below ≈15%, we have excited the fluorophores with a laser power of 1.2 μW. CHO-K1 cells were transfected with the required plasmids and observed after 16 h. Fig 2A shows the brightness values measured for the different FP monomer constructs, either in the cytosol or at the PM. The results follow, as expected, the general trend already described above (Figs 1C and D). No significant difference can be observed between brightness values at this low laser power, with the exception of mp-Gamillus displaying a higher brightness than the other membrane-associated constructs.

Next, we have performed similar experiments on dimer constructs of the same FPs (Fig S2A), in order to calculate the *pf*, as shown in Fig 2B. All *pf* values are in the expected range, around 0.7, and no large difference can be observed between the different fluorophores in general. Noticeably though, mp-Gamillus exhibits *pf* values lower than those of the other FPs (in particular, significantly lower than mp-mGL, ca. 0.6 vs. ca. 0.9).

### Performance comparison of FPs at different pH values

The fluorescence emission intensity of some FPs often decreases at low pH values due to protonation of the chromophore (22, 24, 26). Protonation can induce the transition to dark states and, therefore, negatively affect the accuracy of molecular brightness measurements (28). Here, the pH-dependent fluorescence of the FPs was analyzed via confocal microscopy in the pH range from 5.6 to 9.2 in HEK293T cells using constructs associated to the outer side of the PM via a glycosylphosphatidylinositol (GPI) anchor. The fluorescence emission spectra of the different FPs did not change considerably in the assessed pH range and no photoconversion was observed (Fig S3). Also, the photostability of the FP variants at different pH conditions was qualitatively compared (Fig S4) using an excitation power of 6 μW, on a specific position of the PM, for ca. three minutes. This configuration is usually employed for FCS measurements at the PM and is different from the N&B whole-frame scanning approach used for the experiments described in the previous paragraphs (10). GPI-mEGFP showed a good photostability at all pH conditions, with the fluorescence signal decreasing by only ca. 20% (Figs S4A and S5). GPI-mNG(1x) exhibited fast initial photobleaching, particularly at pH values 5.6 and 9.2, down to ca. 50% of the original signal. At neutral pH, bleaching of GPI-mNG was only slightly higher than that of GPI-mEGFP (Figs S4B and S5). A fast and substantial photobleaching, especially at neutral and low pH was observed for GPI-mGL (Fig S4C and S5). Finally, as shown in Figs S4D and S5, a strong pH-dependency of photostability was observed for GPI-Gamillus: at the highest pH the bleaching was minor (i.e., ca. 20%, similar to GPI-mEGFP), while at neutral and low pH the emission signal dramatically decreased (i.e., to ca. 40% and 10% at pH 7.4 and 5.6, respectively). Overall, mEGFP showed the highest photostability for all pH conditions followed by mNG and then Gamillus (pH < 9.2) and mGL, in agreement with the results from N&B analysis (Fig 1B).

Additionally, the molecular brightness of the different FPs under various pH conditions was evaluated for monomer and dimer membrane-associated GPI-anchored constructs, and the corresponding *pf* values were determined (Figs 3 and S2B). To complement the previous experiments, we used an alternative FFS approach (i.e., sFCS (34)) and an alternative live-cell model (i.e., HEK293T) for this set of experiments. sFCS experiments result in fluorescence auto-correlation curves which can be analyzed to extract parameters such as diffusion coefficients of the labelled proteins, concentration and molecular brightness (see Fig S6 for representative examples). As shown in Fig 3A, the results obtained at pH 7.4 corroborate what was observed also via N&B (Figs 1D and 2A), i.e. that Gamillus overall exhibits a higher brightness compared to the other FPs. This observation holds true for all the tested pH conditions. In general, the observed brightness values did not differ much among each other (being all within a ca. 15% variation interval) and we did not observe an effect of pH, at least at the low laser powers used here.

More marked differences were observed, however, for the *pf* values. In general, this parameter showed a negative correlation with increasing pH for GPI-mEGFP and a positive correlation for GPI-Gamillus. No strong correlation was observed instead for GPI-mGL or GPI-mNG. At pH 5.6, the *pf* of GPI-Gamillus was significantly lower than that of GPI-mEGP and, overall, the lowest among all investigated proteins. At neutral pH, GPI-mGL showed the best performance, in agreement with what we observed for mp-mGL via N&B (Fig 2B). At pH 9.2, no strong differences were observed in general among all FPs, although GPI-Gamillus displayed by trend the highest *pf* value. Notably, we did not observe pH-induced changes in the autocorrelation functions (indicating e.g. significant alterations in the triplet state fraction) for any FP, at this low excitation power (Fig S6). An in-depth study of fluorescence intensity dynamics faster than ca. 1 ms would require, in general, approaches with higher temporal resolution.

## Discussion

PPIs and, specifically, homo-multimerization can be quantified directly in living cells using quantitative fluorescence microscopy approaches based on FFS (e.g., sFCS and N&B) (6–8). The molecular brightness (i.e., photon count rate per molecule) derived from such experiments allows the determination of the oligomeric state of FP-tagged protein complexes. (7, 8, 10). The reliability of such approaches is influenced by the photophysical behavior of the FPs, which can be summed up by the *pf* parameter (7). The *pf* is determined by the specific experimental conditions (e.g. excitation wavelength and power, duration of illumination, geometry of the detection volume), sample environment (pH, ion composition, etc.), intracellular biochemical processes (e.g. maturation time and folding efficiency) and photophysical processes (photobleaching, quantum yield, triplet state formation, protonation- or light-induced “blinking”, long-lived dark states, etc.) (9, 20, 26–28, 34–40). Therefore, three requisites should be ideally met by FPs used in typical FFS studies: i) high photostability, essential to allow time-based measurements under continuous illumination, ii) high molecular brightness, required to achieve sufficient signal-to-noise ratio to detect single molecule fluctuations and iii) a high *pf*, crucial for a large dynamic range and accuracy of oligomerization measurements. In this study, we present a comprehensive analysis of novel GFP variants (mGL and Gamillus) and their suitability for brightness-based oligomerization studies. Both proteins were reported to mature remarkably quickly and to possess a high brightness and medium-low acid sensitivity compared to mEGFP (see Table 1) (21, 24), thus suggesting that they are promising candidates for quantitative FFS studies. These features are, at least partially, shared also by the established GFP variant mNG (23), which is therefore also included in this study.

Previous studies have characterized these FPs and compared them to e.g. mEGFP as standard (see e.g. (41)). Nevertheless, it must be noted that essential parameters, such as brightness, pH sensitivity or photostability, are often measured with different methods and the outcome might depend on the specific setup (23, 24, 41–43). Thus, it is reasonable to compare the FPs systematically using specific conditions and approaches resembling those of actual FFS experiments. Following this logic, we used typical FFS approaches (i.e., N&B and sFCS) to quantify the stability, brightness, pH sensitivity and *pf* of mEGFP, mNG, Gamillus and mNG.

A first unexpected result is that, in general, we do not observe strikingly different brightness values between each FP. While this observation can likely be explained by the distinctive experimental setup (e.g. very low excitation power, longer acquisition times, specific intra-cellular localization (21, 42, 44)), it is nevertheless relevant for users planning similar FFS-based investigations. Furthermore, the brightness measured via FFS is an average value derived from a statistical analysis of single molecule fluctuations. Such analysis is different from bulk methods that measure the fluorescence emission in a whole cell as a readout of brightness. While remaining both valid, the different approaches might provide information about different aspects of the fluorescent system. Finally, in our experimental conditions, Gamillus displayed a higher brightness compared to the other FPs, at least when located at the PM or while using higher laser powers. This observation was independently confirmed via both sFCS and N&B analysis.

Both techniques also indicate that the higher brightness of Gamillus is counterbalanced by a stronger tendency to photobleach, at least in the conditions used in our experiments (e.g., pH 7.4). A similarly low photostability was observed also for mGL, while mNG and mEGFP proved to be remarkably stable under continuous irradiation. Of interest, the contrast with reports indicating a strong photostability of mGL and Gamillus might be only apparent. A general comparison of these observations with previous studies is complicated by the fact that, often, different conditions and experimental setups result in different apparent photochemical behavior. For example, mEGFP was shown to be less stable than mNG under widefield illumination, more stable under laser illumination (23) and similarly stable using a spinning disk confocal microscope (42). Nevertheless, the photostability observed here at pH 7.4 for mEGFP is in the same range as previously reported (23, 45). Also, mGL was reported to be less photostable than Clover that, in turn, is less photostable than mEGFP (21, 45), in agreement with our observations. On the other hand, in contrast to our observation, Shinoda *et al*. showed that Gamillus has a 2-fold higher photostability than mEGFP under widefield illumination and acidic conditions (24). Once more, a possible explanation for such discrepancies might reside in the considerably different experimental conditions which, in this case, might be specifically relevant for FFS-based multimerization studies.

Next, we have measured the *pf* values for the different FPs, since this is a parameter of fundamental importance for quantitative multimerization studies. Our results indicate that, in general, all the examined FPs perform relatively well (i.e., *pf* ca. 0.7) and definitely better than most red or blue FPs (7, 19). In particular, mGL excels in this context, with a *pf* value of 0.8 and above, as confirmed by both N&B and sFCS in different cell models. This might be due to its fast maturation rate (21).

In the second part of our work, we compared the FPs under different pH conditions. This is relevant for live-cell studies with FPs targeting e.g. acidic organelles, such as endosomes, secretory granules, lysosomes, and Golgi-Network (pH range ca. 4.7-8 (46)), which play a role for the sorting, transport and degradation of proteins (24, 25). In general, mNG and mGL did not exhibit a special sensitivity to changes in pH values between 5.6 and 9.2. Neither of the monitored parameters (photostability, brightness or *pf*)displayed a significant correlation with pH values.

In contrast, mEGFP showed an increase of *pf* at acidic pH, with this value being significantly higher than that of Gamillus. We did not observe a strong decrease in mEGFP brightness at low pH values, as it was instead previously reported (26, 28, 43, 44). This might be due to the low laser powers employed in our studies, since a control experiment performed at a ca. four-fold higher laser power resulted in a decrease in mEGFP brightness of ca. 40% (data not shown). Apart from light-induced photochemical effect, it might also be possible that the data spread obtained at lower excitation power could partially mask a decrease in brightness. In any case, our data show that the brightness of mEGFP at pH 5.6 is indeed slightly lower than that of the other FPs.

Also Gamillus was influenced by changes in pH value. First, we observed that its brightness remained higher than that of mEGFP, especially at low pH values. Interestingly, the photostability and the *pf* decreased dramatically as a function of decreasing pH values. In contrast with the idea of Gamillus being a FP particularly useful under acidic conditions (24), our data indicate that this fluorophore might perform best in basic environments in the context of FFS measurements. It is possible that strong differences between Gamillus and, e.g., mEGFP might be observed instead at pH values below those explored in this work (i.e. pH <5.5).

In conclusion, our results indicate that, in general, mEGFP is a very good choice for the quantification of multimerization via FFS, also at pH values between 5.6 and 7.4. Although its brightness is lower than other examined FPs, its remarkable photostability would allow using higher excitation powers (thus increasing its brightness). Bright FPs, such as Gamillus, could be efficiently used for e.g. qualitative imaging with short acquisition times and low excitation powers. At neutral pH conditions, a remarkably high *pf* was observed for mGL, although care should be taken in experiments in which photobleaching might represent an issue. Finally, at basic pH conditions (e.g., up to 9.2), Gamillus represents an optimal choice based on a good photostability, high *pf* and brightness.

## Supporting information

Data for Figs 1, 2, 3, S2, S3, S5

## Abbreviations

FFS: fluorescence fluctuation spectroscopy
FP: fluorescence protein
N&B: number and brightness
*pf*: fluorescence probability
PM: plasma membrane
PPI: protein-protein-interaction
sFCS: scanning fluorescence correlation spectroscopy

## Author Contribution

Research Planning, A.P., A.K.A. and S.C; Investigation, A.P. and A.K.A.; Data Analysis, A.P., A.K.A. and S.C.; Writing – Original Draft Preparation, A.P.; Writing – Review & Editing, A.K.A., V.D., and S.C.; Software, V.D., A.P. and S.C.; Supervision, S.C.

## Acknowledgements

We thank all the members of the Physical Biochemistry group for useful feedback. VD is grateful for support by an HFSP long-term postdoctoral fellowship (HFSP LT0058/2022-L). This work was partially supported by the Deutsche Forschungsgemeinschaft (DFG) project number 407961559 to S.C.

## Supporting Information

### Supporting Materials and Methods

**Table S1: Overview of the linker sequences of the homo-dimers for the different fluorescence protein (FP) constructs used in this study.** mp: myristoylated and palmitoylated peptide, GPI: glycosylphosphatidylinositol anchor.

**Table S2: Overview of molecular brightness and bleached fraction values measured via number and brightness (N&B) analysis for all FPs in the cytosol and at the plasma membrane (PM).** Data correspond to Fig 1 of the main manuscript.

**Fig S1: Schematic overview of the experimental setup.** (A) Overview of the different fluorescent proteins (FPs) used in this study: cytosolic/soluble FPs, membrane associated constructs with the myristoylated and palmitoylated (mp) peptide and glycosylphosphatidylinositol (GPI)-anchor linked to the FP as a monomer or homo-dimer. The FPs used in this study were mEGFP, mNeonGreen (mNG), mGreenLantern (mGL) and Gamillus. (B) Overview of the experimental procedure. One day after seeding, cells are transfected. On the day after, cells are directly used for Number and Brightness (N&B) measurements or are washed and equilibrated in HEPES-Buffer with the appropriate pH before each measurement for scanning fluorescence correlation spectroscopy (sFCS) and spectral analysis. (C) N&B acquisition results in a three-dimensional (x-y-time) image stack. A ROI is selected around a cell or membrane region. Then, brightness (ε) values are calculated in each pixel. The results are then visualized as average intensity (<I>) and average brightness (<ε>) map and histogram. (D) sFCS measurements are performed perpendicular to the plasma membrane (PM), as also shown in panel (B). Scan lines (represented as kymographs) are aligned and intensity at PM is integrated. Then, the brightness and the ACF are calculated from the intensity trace and the ACF is analysed with a two-dimensional diffusion model.

**Fig S2: Brightness comparison for different green FP homo-dimers.** (A): N&B measurements were performed ≈16 h after transfection in CHO-K1 cells, using a laser power of 1.2 μW. Box plot of the molecular brightness in kHz for the examined cytosolic and membrane-anchored FP (i.e., mp-FP) homo-dimers. Each point represents the average value measured in a single cell, pooled from three independent experiments. (B): sFCS measurements were performed in HEK293T cells for different pH conditions (pH 5.6, pH 7.4, and pH 9.2) ≈16 h after transfection, with a laser power of 1.5 μW. Box plots with single data points from three independent experiments show the molecular brightness in kHz for the homo-dimers of GPI-mEGFP, GPI-mNG, GPI-mGL and GPI-Gamillus. Median values and whiskers ranging from minimum to maximum values are displayed. Sample size, mean, median, and interquartile range (IQR) are indicated in the graph. Statistical significance was determined for both plots using one-way ANOVA Tukey’s multiple comparison test; * p < 0.05, ** p < 0.005, ***p < 0.0005, ****p < 0.0001.

**Fig S3: Normalized FP emission spectra at different pH values.** Average emission spectra of GPI-mEGFP (A), GPI-mNG (B), GPI-mGL (C), and GPI-Gamillus (D) measured via spectral imaging (23 spectral channels from 491 nm to 695 nm) using 488 nm excitation on HEK 293T cells supplemented with buffer at different pH values (5.6, 7.4 and 9.2). At each pH value, ca. 10 cells were imaged, acquiring ten frames. To obtain the average emission spectra, pixels corresponding to the PM were semi-manually segmented (manual selection followed by removal of pixels with intensities below 25% of the maximum pixel intensity in the selected region) and detected spectra averaged over all pixels and cells measured at each pH.

**Fig S4: Comparison of photostability for different monomeric green FPs under different pH conditions via sFCS measurements.** HEK293T cells were transfected with the appropriate FP construct, washed on the next day with HEPES buffer with the corresponding pH (pH 5.2 (magenta), pH 7.4 (green), and pH 9.2 (blue)) and then measured with a laser power of 6 μW, which is 4-fold higher than that used for standard sFCS measurements. The emission intensities were normalized to the initial values. Representative photobleaching curves are shown for GPI-mEGFP (A), GPI-mNG (B), GPI-mGL (C) and GPI-Gamillus (D). Solid line represents a double exponential fit, as guide to the eye.

**Fig S5: Comparison of bleached fractions for the examined monomeric FPs, at different pH values**. The decrease of the fluorescence signal has been quantified after a 180 s sFCS measurement, as exemplified in Fig S4. For each FP, between 13 and 15 measurements from 3 independent samples were performed. Statistical significance was determined using one-way ANOVA Tukey’s multiple comparison test; * p < 0.05, ** p < 0.005, ***p < 0.0005. Different symbols refer to comparisons to GPI-mEGFP (*), GPI-mNG (#), GPI-mGL (†).

**Fig S6: Representative sFCS autocorrelation functions and fit curves obtained for cells expressing GPI-FPs, under different pH conditions.** Fit curves (solid line) were obtained by fitting a two-dimensional diffusion model to the data, as described in the main text (Methods section).

