## Supplementary Information for "Benchmarking of novel green fluorescent proteins for the quantification of protein oligomerization in living cells"

#### Supporting Materials and Methods

##### Fluorescent protein constructs

For the cloning of all constructs, standard PCRs with custom-designed primers were performed, followed by digestion with FastDigest restriction enzymes and ligation with T4-DNA-Ligase according to the manufacturer's instructions. All enzymes and reagents were purchased from Thermo Fisher Scientific (Waltham, MA, USA) and primers were acquired from Sigma Aldrich trademark of Merck KGaA (Darmstadt, Germany). Each construct was verified by Sanger sequencing (LGC Genomics GmbH, Berlin, Germany).

The plasmids encoding the monomeric and dimeric fluorescence protein (FP) mEGFP for i) cytoplasmic expression (mEGFP(1x) and mEGFP(2x)), ii) inner leaflet plasma membrane (PM) localization (FP linked to a myristoylated and palmitoylated (mp) peptide, mp-mEGFP(1x) and mp-mEGFP(2x)) and iii) outer leaflet PM localization (FP linked to a glycosylphosphatidylinositol (GPI)-anchor, GPI-mEGFP(1x)) were previously described [1, 2]. GPI-mEGFP(2x) was cloned by amplifying the GPI signal peptide ($GPI_{sp}$) linked to mEGFP from the GPI-mEGFP(1x) plasmid and inserting it into mp-EGFP(1x), using digestion with HindIII and NotI. Afterwards, the GPI-anchor C-terminally linked to mEGFP was amplified from GPI-mEGFP(1x) and cloned into the $GPI_{sp}$-mEGFP construct, using digestion with MluI and NotI.

The plasmids encoding the monomeric and dimeric FP mGreenLantern (mGL) for i) cytoplasmic expression (mGL(1x) and mGL (2x)), ii) inner leaflet PM localization (FP linked to a mp peptide, mp-mGL(1x) and mp-mGL(2x)) and iii) outer leaflet PM localization (FP linked to a GPI-anchor, GPI-mGL(1x) and GPI-mGL(2x)) were created by cloning of the monomeric mGL cassette from mGreenLantern_pcDNA3.1, a gift from Benjamin C. Campell (Helen and Robert Appel Alzheimer's Disease Research Institute, Weill Cornell Medicine, New York; Addgene plasmid #161912). To obtain all constructs in the same vector backbone, mGL was amplified from the mGreenLantern_pcDNA3.1 vector with and without a N-terminal linked mp peptide and cloned into the mEGFP-C1 (gift from Michael Davidson, Addgene plasmid #54759) by digestion with AgeI and Kpn21 or NheI and Kpn21, respectively. Subsequently, an additional amplified monomeric cassette from the mGreenLantern_pcDNA3.1 vector was ligated into mGL(1x) and mp-mGL(1x) by digestion with BglII and Kpn21 to generate mGL(2x) and mp-mGL(2x). Next, mp-mGL(1x) was digested with AgeI and BsrGI and cloned into GPI-mEGFP(1x) to generate GPI-mGL(1x). To generate the GPI dimer construct, mGL from the GPI-mGL(1x) construct was ligated into the $GPI_{sp}$-mEGFP plasmid by digestion with HindIII and BsrGI to generate the $GPI_{sp}$-mGL construct. Finally, the GPI-anchor C-terminally linked to mGL was amplified from the GPI-mGL(1x) plasmid and cloned into the $GPI_{sp}$-mGL construct, using digestion with MluI and NotI.

The plasmids encoding the monomeric and dimeric FP mNeonGreen (mNG) for i) cytoplasmic expression (mNG(1x) and mNG(2x)), ii) inner leaflet PM localization (FP linked to a mp peptide, mp-mNG (1x) and mp-mNG (2x)) and iii) outer leaflet PM localization (FP linked to a GPI-anchor, GPI-mNG(1x) and GPI-mNG(2x)) were created by cloning of the monomeric mNG cassette from pHAGE mNeonGreen-Core (HBc from HBV) IRES puro, a gift from Raphael Gaudin (Addgene plasmid #122202). To have all constructs in the same vector backbone, mNG was amplified from the pHAGE mNeonGreen-Core (HBc from HBV) IRES puro vector and cloned into mEGFP-C1 (gift from Michael Davidson, Addgene plasmid #54759) by digestion with AgeI and Kpn21 to generate mNG(1x). To obtain mNG(2x), mNG was amplified from mNG(1x) and the PCR product inserted into mNG(1x) by digestion with KpnI and EcoRI. Mp-mNG(1x) was generated by a ligation of the digest product of mNG(1x) with AgeI and BsrGI into the mp-mEGFP(1x) plasmid. Afterwards, the monomeric mNG cassette was amplified from the mp-NG(1x) construct and cloned into the mNG(2x) by digestion with NheI and BsrGI to obtain the mp-mNG(2x) plasmid. To generate GPI-mNG(1x), mp-mNG(1x) was digested with AgeI and BsrGI and cloned into GPI-mEGFP(1x) to generate GPI-mNG(1x). To obtain GPI-mNG(2x), mNG from the GPI-mNG(1x) construct was ligated into the $GPI_{sp}$-mEGFP plasmid after digestion with HindIII and BsrGI to generate the $GPI_{sp}$-mNG construct. Subsequently, the GPI-anchor C-terminal linked to mNG was amplified from the GPI-mNG(1x) plasmid and cloned into the $GPI_{sp}$-mNG construct, using digestion with MluI and NotI.

The plasmids encoding the monomeric and dimeric FP Gamillus for i) cytoplasmic expression (Gamillus(1x) and Gamillus(2x)), ii) inner leaflet PM localization (FP linked to a mp peptide, mp-Gamillus(1x) and mp-Gamillus(2x)) and iii) outer leaflet PM localization (FP linked to a GPI-anchor, GPI-Gamillus(1x) and GPI-Gamillus(2x)) were created by cloning of the monomeric Gamillus cassette from Gamillus/pcDNA3, a gift from Takeharu Nagai (Addgene plasmid #124837). To obtain all constructs in the same vector backbone, Gamillus was amplified from Gamillus/pcDNA3 and inserted into mEGFP-C1 (gift from Michael Davidson, Addgene plasmid #54759) by digestion with AgeI and Kpn21 to generate Gamillus(1x). To obtain Gamillus(2x), Gamillus was additionally amplified from Gamillus/pcDNA3 and cloned into Gamillus(1x) by digestion with Kpn21 and BglIII. The mp-Gamillus(1x) construct was generated by cloning the monomeric Gamillus cassette from Gamillus(1x) into mp-mEGFP(1x) after digestion with AgeI and BsrGI. The mp-Gamillus(2x) construct was obtained by a stepwise cloning. First, monomeric Gamillus cassette was amplified from Gamillus/pcDNA3 and cloned into mp-mNG(2x) by digestion with AgeI and BglII. Finally, the Gamillus cassette was additionally amplified from the Gamillus/pcDNA3 vector and inserted into the mp-Gamillus-mNG construct after digestion with BglII and EcoRI. Next, mp-mGamillus(1x) was digested with AgeI and BsrGI and cloned into GPI-mEGFP(1x) to generate GPI-mGamillus(1x). To generate the GPI dimer construct, the Gamillus cassette from the GPI-Gamillus(1x) construct was ligated into the $GPI_{sp}$-mEGFP plasmid by digestion with HindIII and BsrGI to generate the $GPI_{sp}$-Gamillus construct. Subsequently, the GPI-anchor C-terminal linked to mGamillus was amplified from the GPI-Gamillus(1x) plasmid and cloned into the $GPI_{sp}$-Gamillus construct, using digestion with MluI and NotI.

An overview of the linker sequences between the FPs is provided in Table S1.

### Supporting Tables

**Table S1: Overview of the linker sequences of the homo-dimers for the different fluorescence protein (FP) constructs used in this study.** mp: myristoylated and palmitoylated peptide, GPI: glycosylphosphatidylinositol anchor.

| Construct | Linker length between FPs | Linker sequence between FPs |
|---|---|---|
| **mEGFP(2x)** | 7 aa | SGLRSRG |
| **mp-mEGFP(2x)** | 7 aa | SGLRSRG |
| **GPI-mEGFP(2x)** | 12 aa | CTSRLPLHLRNA |
| **mGL(2x)** | 10 aa | SGPPAAAPPV |
| **mp-mGL(2x)** | 10 aa | SGPPAAAPPV |
| **GPI-mGL(2x)** | 10 aa | PPAAAPPERV |
| **mNG(2x)** | 10 aa | SGLRSRAQAS |
| **mp-mNG(2x)** | 7 aa | RSRAQAS |
| **GPI-mNG(2x)** | 10 aa | PPAAAPPERV |
| **Gamillus(2x)** | 10 aa | SGPPAAAPPV |
| **mp-Gamillus(2x)** | 10 aa | RSPPAAAPPV |
| **GPI-Gamillus(2x)** | 10 aa | PPAAAPPERV |

**Table S2: Overview of molecular brightness and bleached fraction values measured via number and brightness (N&B) analysis for monomeric FPs in the cytosol and at the plasma membrane (PM), for different laser powers.** Data correspond to Fig 1 of the main manuscript.

| Construct | Laser power | Bleached fraction | | Brightness [kHz] | | |
|---|---|---|---|---|---|---|
| | [µW] | mean | SEM | mean | SEM | n |
| mEGFP(1x) | 0.6 | 0.020 | 0.009 | 0.69 | 0.23 | 3 |
| | 1.2 | 0.023 | 0.004 | 0.89 | 0.06 | 28 |
| | 2.4 | 0.036 | 0.008 | 1.23 | 0.08 | 21 |
| mNG(1x) | 0.6 | 0.024 | 0.013 | 0.42 | 0.07 | 7 |
| | 1.2 | 0.054 | 0.014 | 0.95 | 0.05 | 28 |
| | 2.4 | 0.036 | 0.007 | 1.73 | 0.10 | 24 |
| mGL(1x) | 0.6 | 0.027 | 0.011 | 0.36 | 0.16 | 3 |
| | 1.2 | 0.117 | 0.007 | 0.82 | 0.04 | 25 |
| | 2.4 | 0.300 | 0.025 | 2.03 | 0.10 | 26 |
| Gamillus(1x) | 0.6 | 0.051 | 0.010 | 0.27 | 0.07 | 8 |
| | 1.2 | 0.116 | 0.010 | 0.95 | 0.08 | 33 |
| | 2.4 | 0.173 | 0.018 | 2.15 | 0.14 | 32 |
| mp-mEGFP(1x) | 0.6 | 0.051 | 0.022 | 1.30 | 0.40 | 4 |
| | 1.2 | 0.049 | 0.013 | 1.36 | 0.13 | 14 |
| | 2.4 | 0.041 | 0.007 | 2.82 | 0.23 | 29 |
| mp-mNG(1x) | 0.6 | 0.030 | 0.008 | 1.20 | 0.50 | 6 |
| | 1.2 | 0.036 | 0.006 | 1.23 | 0.11 | 17 |
| | 2.4 | 0.037 | 0.005 | 2.52 | 0.22 | 30 |
| mp-mGL(1x) | 0.6 | 0.056 | 0.009 | 0.73 | 0.23 | 6 |
| | 1.2 | 0.138 | 0.016 | 1.43 | 0.14 | 22 |
| | 2.4 | 0.315 | 0.025 | 2.37 | 0.25 | 8 |
| mp-Gamillus(1x) | 0.6 | 0.042 | 0.014 | 0.92 | 0.27 | 8 |
| | 1.2 | 0.146 | 0.019 | 1.69 | 0.17 | 22 |
| | 2.4 | 0.232 | 0.020 | 3.60 | 0.30 | 37 |

#### Supporting Figures

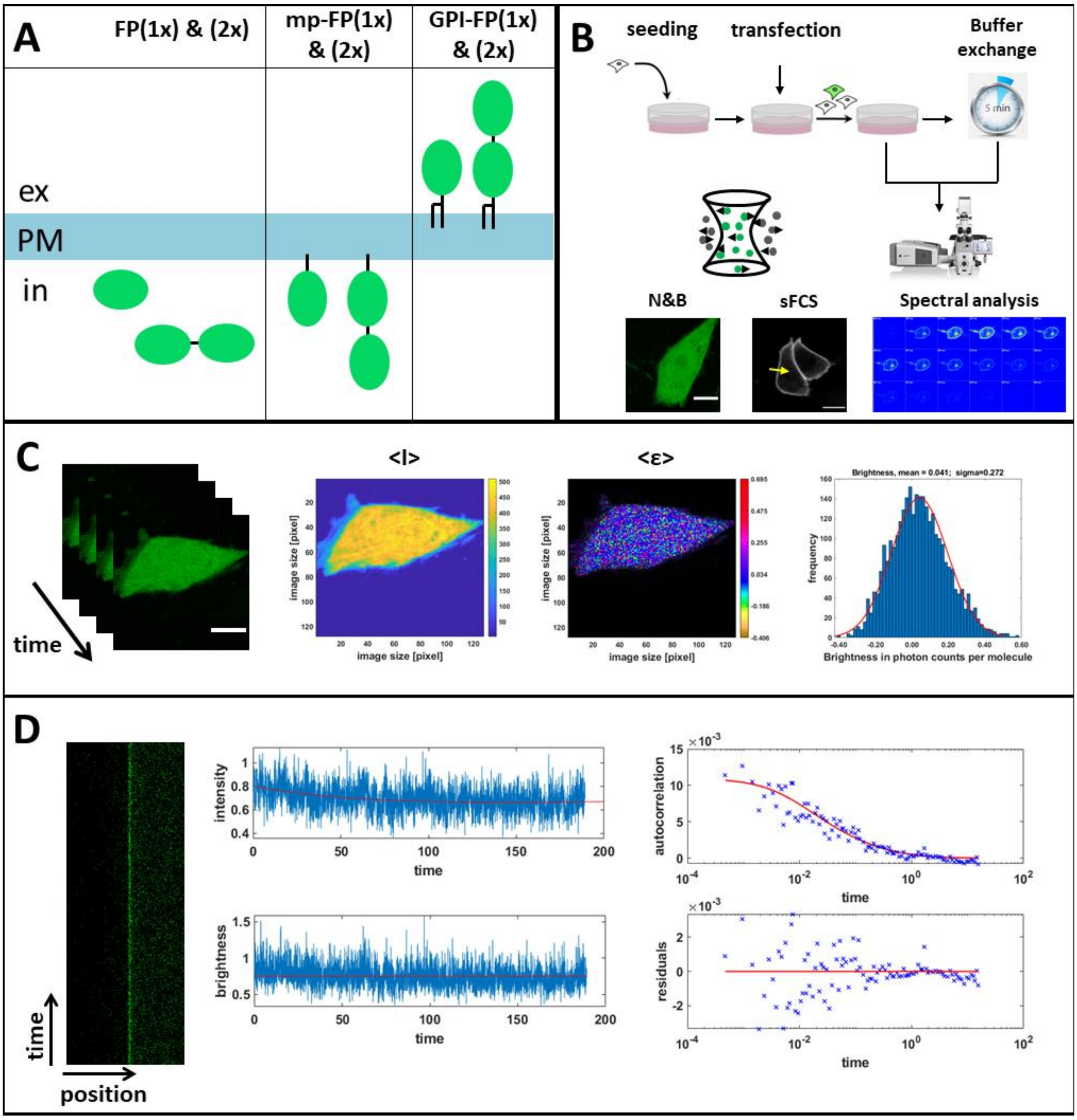


**Fig S1: Schematic overview of the experimental setup.** (A) Overview of the different fluorescent proteins (FPs) used in this study: cytosolic/soluble FPs, membrane associated constructs with the myristoylated and palmitoylated (mp) peptide and glycosylphosphatidylinositol (GPI)-anchor linked to the FP as a monomer or homo-dimer. The FPs used in this study were mEGFP, mNeonGreen (mNG), mGreenLantern (mGL) and Gamillus. (B) Overview of the experimental procedure. One day after seeding, cells are transfected. On the day after, cells are directly used for Number and Brightness (N&B) measurements or are washed and equilibrated in HEPES-Buffer with the appropriate pH before each measurement for scanning fluorescence correlation spectroscopy (sFCS) and spectral analysis. (C) N&B acquisition results in a three-dimensional (x-y-time) image stack. A ROI is selected around a cell or membrane region. Then, brightness (ε) values are calculated in each pixel. The results are then visualized as average intensity (<I>) and average brightness (<ε>) map and histogram. (D) sFCS measurements are performed perpendicular to the plasma membrane (PM), as also shown in panel (B). Scan lines (represented as kymographs) are aligned and intensity at PM is integrated. Then, the brightness and the ACF are calculated from the intensity trace and the ACF is analyzed with a two-dimensional diffusion model.

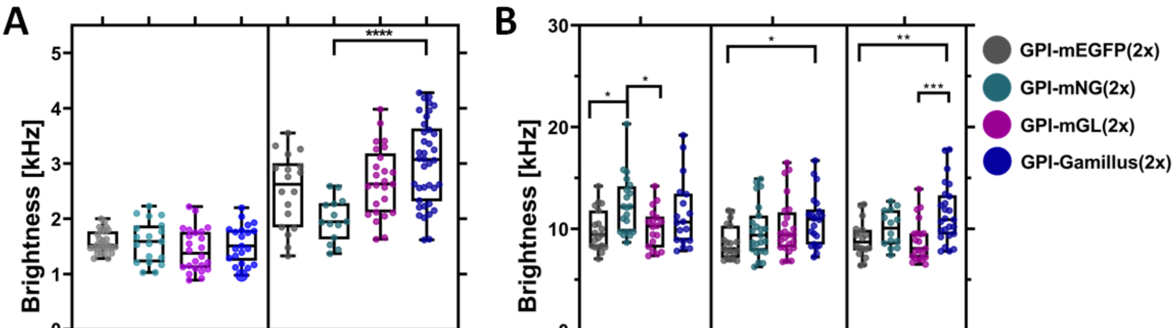


| | mEGFP(2x) | mNG(2x) | mGL(2x) | Gamillus(2x) | mp-mEGFP(2x) | mp-mNG(2x) | mp-mGL(2x) | mp-Gamillus(2x) |
|---|---|---|---|---|---|---|---|---|
| n | 30 | 19 | 24 | 21 | 18 | 14 | 25 | 38 |
| mean | 1.6 | 1.6 | 1.4 | 1.6 | 2.5 | 2.0 | 2.7 | 3.0 |
| median | 1.5 | 1.6 | 1.4 | 1.5 | 2.6 | 2.0 | 2.6 | 3.1 |
| IQR | 0.3 | 0.7 | 0.7 | 0.6 | 1.2 | 0.6 | 1.1 | 1.3 |

| | pH 5.6 | | | | pH 7.4 | | | | pH 9.2 | | | |
|---|---|---|---|---|---|---|---|---|---|---|---|---|
| n | 19 | 20 | 18 | 18 | 19 | 23 | 24 | 22 | 18 | 14 | 21 | 25 |
| mean | 9.9 | 12.4 | 10.0 | 11.5 | 8.6 | 9.8 | 10.1 | 10.8 | 9.0 | 10.0 | 8.8 | 11.5 |
| median | 9.4 | 12.2 | 10.3 | 10.7 | 8.0 | 9.4 | 9.4 | 11.0 | 8.7 | 10.1 | 8.1 | 10.9 |
| IQR | 3.6 | 4.5 | 3.1 | 4.7 | 3.2 | 3.5 | 3.4 | 3.5 | 2.0 | 3.4 | 2.4 | 3.9 |

**Fig S2: Brightness comparison for different green FP homo-dimers.** (A): N&B measurements were performed ≈16 h after transfection in CHO-K1 cells, using a laser power of 1.2 µW. Box plot of the molecular brightness in kHz for the examined cytosolic and membrane-anchored FP (i.e., mp-FP) homo-dimers. Each point represents the average value measured in a single cell, pooled from three independent experiments. (B): sFCS measurements were performed in HEK293T cells for different pH conditions (pH 5.6, pH 7.4, and pH 9.2) ≈16 h after transfection, with a laser power of 1.5 µW. Box plots with single data points from three independent experiments show the molecular brightness in kHz for the homo-dimers of GPI-mEGFP , GPI-mNG , GPI-mGL and GPI-Gamillus. Median values and whiskers ranging from minimum to maximum values are displayed. Sample size, mean, median, and interquartile range (IQR) are indicated in the graph. Statistical significance was determined for both plots using one-way ANOVA Tukey´s multiple comparison test; * $p < 0.05$, ** $p < 0.005$, ***$p < 0.0005$, ****$p < 0.0001$.

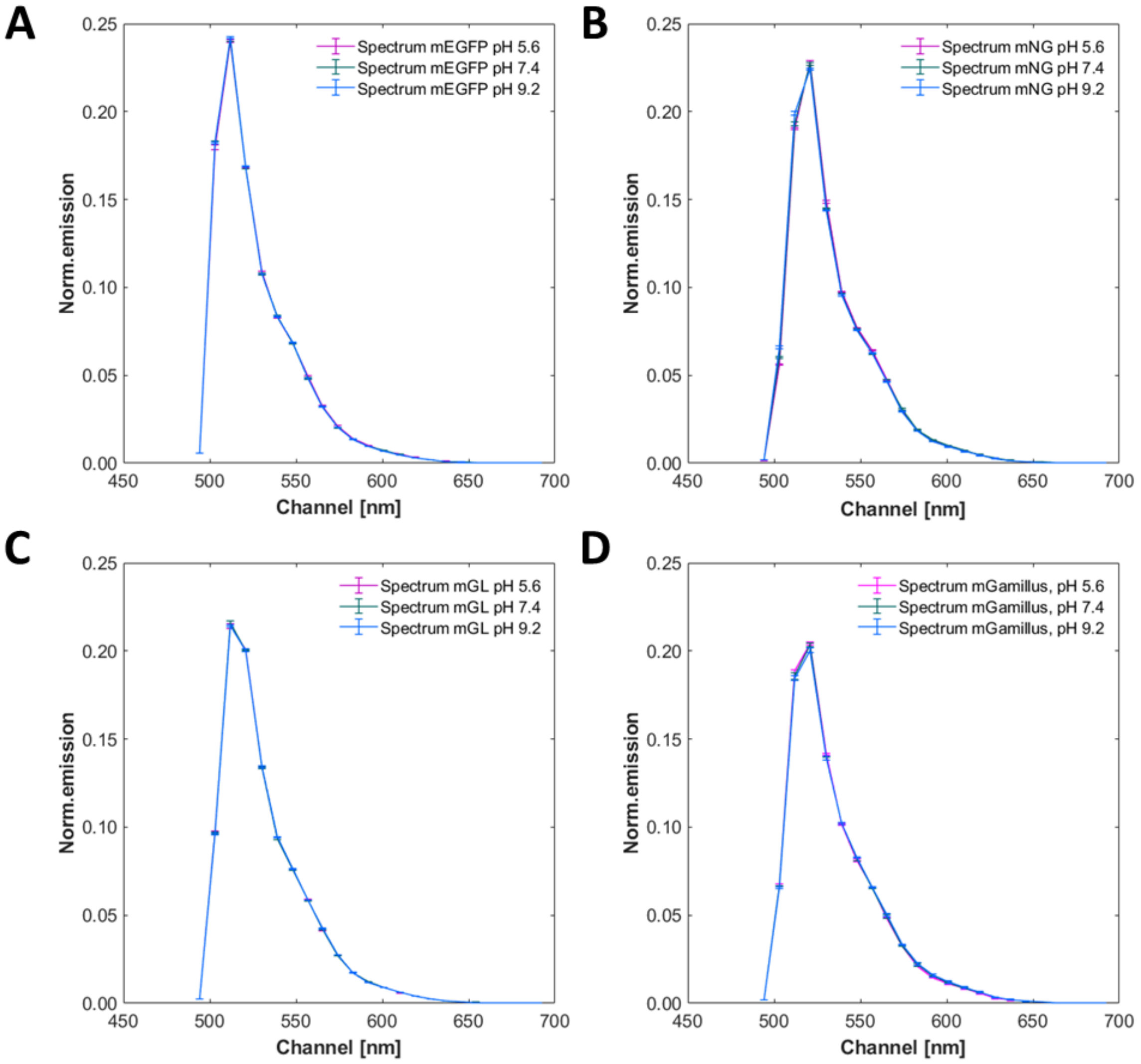


**Fig S3: Normalized FP emission spectra at different pH values.** Average emission spectra of GPI-mEGFP (A), GPI-mNG (B), GPI-mGL (C), and GPI-Gamillus (D) measured via spectral imaging (23 spectral channels from 491 nm to 695 nm) using 488 nm excitation on HEK 293T cells supplemented with buffer at different pH values (5.6, 7.4 and 9.2). At each pH value, ca. 10 cells were imaged, acquiring ten frames. To obtain the average emission spectra, pixels corresponding to the PM were semi-manually segmented (manual selection followed by removal of pixels with intensities below 25% of the maximum pixel intensity in the selected region) and detected spectra averaged over all pixels and cells measured at each pH.

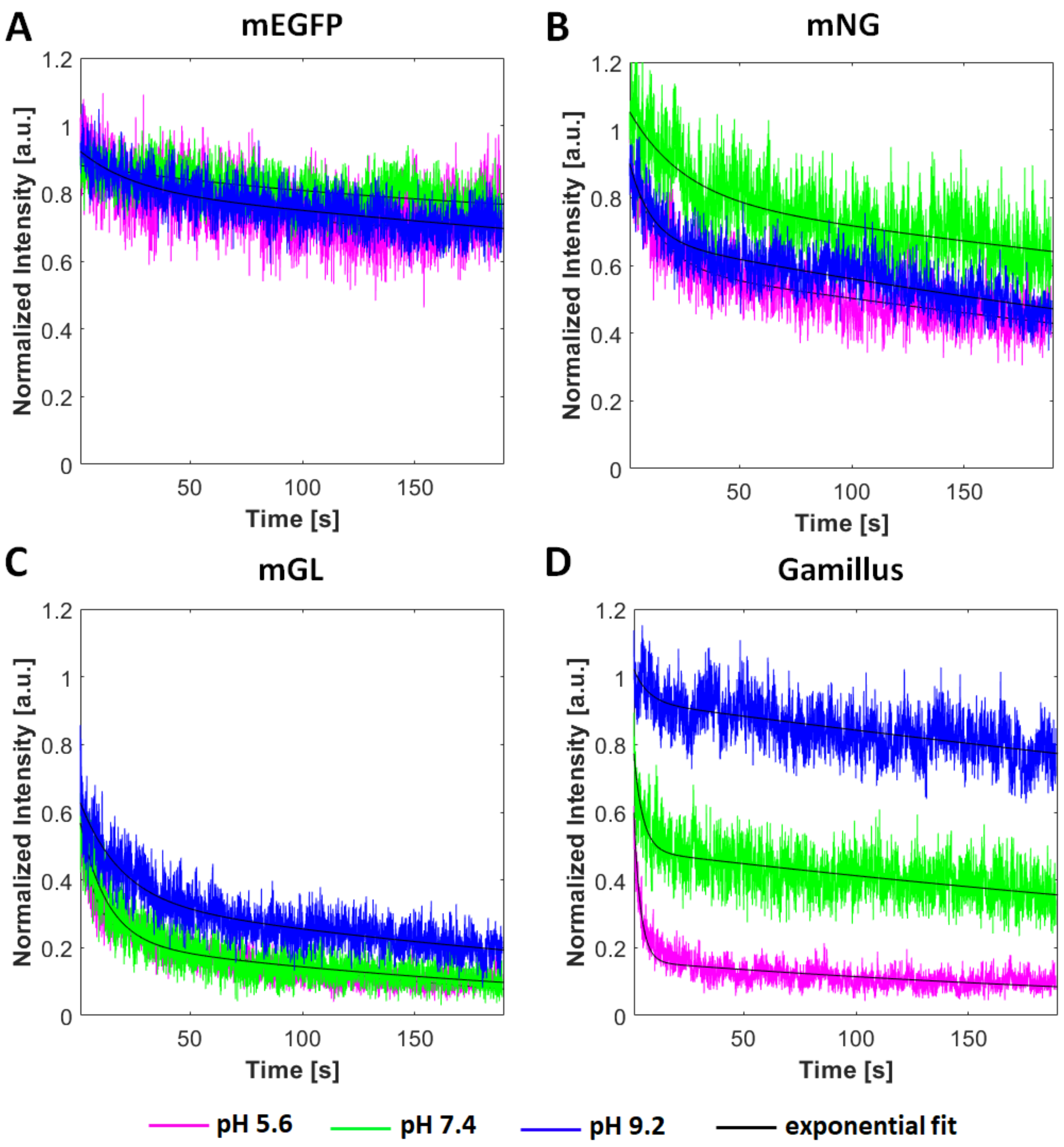


**Fig S4: Comparison of photostability for different monomeric green FPs under different pH conditions via sFCS measurements.** HEK293T cells were transfected with the appropriate FP construct, washed on the next day with HEPES buffer with the corresponding pH (pH 5.2 (magenta), pH 7.4 (green), and pH 9.2 (blue)) and then measured with a laser power of 6 µW, which is 4-fold higher than that used for standard sFCS measurements. The emission intensities were normalized to the initial values. Representative photobleaching curves are shown for GPI-mEGFP (A), GPI-mNG (B), GPI-mGL (C) and GPI-Gamillus (D). Solid line represents a double exponential fit, as guide to the eye.

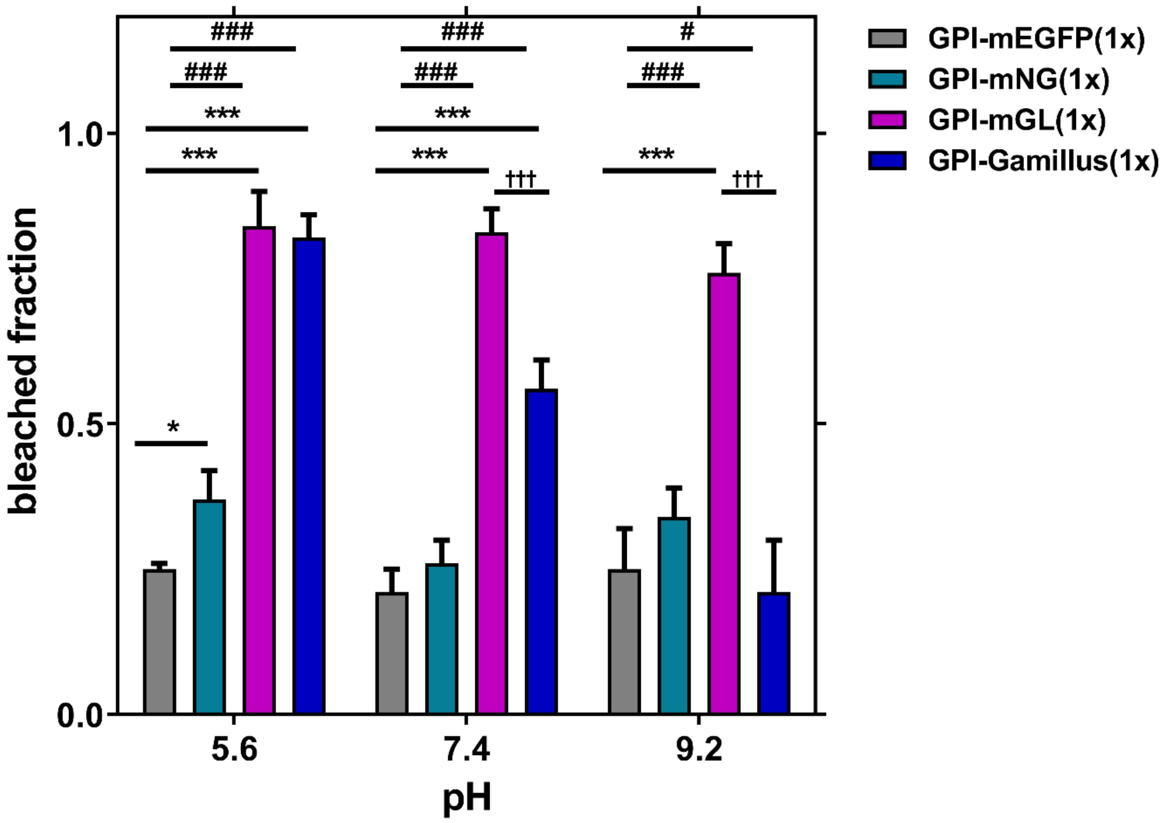


**Fig S5: Comparison of bleached fractions for the examined monomeric FPs, at different pH values**. The decrease of the fluorescence signal has been quantified after a 180 s sFCS measurement, as exemplified in Fig S4. For each FP, between 12 and 15 measurements from 3 independent samples were performed. Statistical significance was determined using one-way ANOVA Tukey´s multiple comparison test; * $p < 0.05$, ** $p < 0.005$, ***$p < 0.0005$. Different symbols refer to comparisons to GPI-mEGFP (*), GPI-mNG (#), GPI-mGL (†).

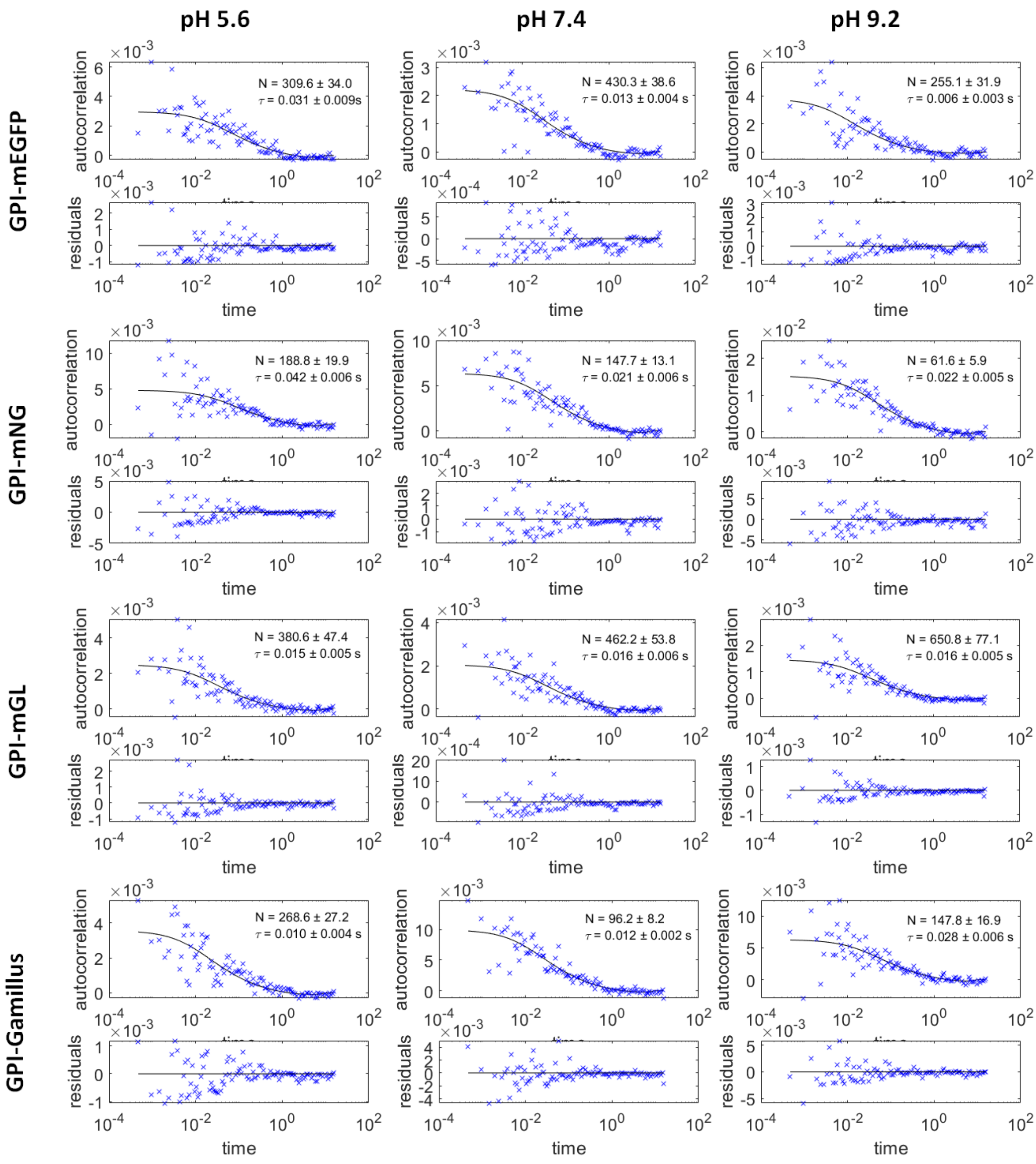


**Fig S6: Representative sFCS autocorrelation functions and fit curves obtained for cells expressing GPI-FPs, under different pH conditions.** Fit curves (solid line) were obtained by fitting a two-dimensional diffusion model to the data, as described in the main text (Methods section).
